# Genetic association study of psychotic experiences in UK Biobank

**DOI:** 10.1101/583468

**Authors:** Sophie E. Legge, Hannah J. Jones, Kimberley M. Kendall, Antonio F. Pardiñas, Georgina Menzies, Mathew Bracher-Smith, Valentina Escott-Price, Elliott Rees, Katrina A.S. Davis, Matthew Hotopf, Jeanne E. Savage, Danielle Posthuma, Peter Holmans, George Kirov, Michael J. Owen, Michael C. O’Donovan, Stanley Zammit, James T.R. Walters

**Author notes:** Both authors contributed equally to this project (Stanley Zammit, James T.R. Walters). Corresponding author*: Professor James Walters, MRC Centre for Neuropsychiatric Genetics and Genomics, Division of Psychological Medicine and Clinical Neurosciences, School of Medicine, Cardiff University, Cardiff, UK.

## Abstract

Psychotic experiences, such as hallucinations and delusions, are reported by approximately 5%-10% of the general population, though only a small proportion of individuals develop psychotic disorders such as schizophrenia or bipolar disorder. Studying the genetic aetiology of psychotic experiences in the general population, and its relationship with the genetic aetiology of other disorders, may increase our understanding of their pathological significance. Using the population-based UK Biobank sample, we performed the largest genetic association study of psychotic experiences in individuals without a psychotic disorder. We conducted three genome-wide association studies (GWAS) for (i) any psychotic experience (6123 cases vs. 121,843 controls), (ii) distressing psychotic experiences (2143 cases vs. 121,843 controls), and (iii) multiple occurrence psychotic experiences (3337 cases vs. 121,843 controls). Analyses of polygenic risk scores (PRS), genetic correlation, and copy number variation (CNV) were conducted to assess whether genetic liability to psychotic experiences is shared with schizophrenia and/or other neuropsychiatric disorders and traits. GWAS analyses identified four loci associated with psychotic experiences including a locus in Ankyrin-3 (*ANK3*, OR=1.16, *p*=3.06 × 10^−8^) with any psychotic experience and a locus in cannabinoid receptor 2 gene (*CNR2,* OR=0.66, *p*=3.78×10^−8^) with distressing psychotic experiences. PRS analyses identified associations between psychotic experiences and genetic liability for schizophrenia, major depressive disorder, and bipolar disorder, and these associations were stronger for distressing psychotic experiences. Genetic correlation analysis identified significant genetic correlations between psychotic experiences and major depressive disorder, schizophrenia, autism spectrum disorder and a cross-disorder GWAS. Individuals reporting psychotic experiences had an increased burden of CNVs previously associated with schizophrenia (OR=2.04, *p*=2.49×10^−4^) and of those associated with neurodevelopmental disorders more widely (OR=1.75, *p*=1.41×10^−3^). In conclusion, we identified four genome-wide significant loci in the largest GWAS of psychotic experiences from the population-based UK Biobank sample and found support for a shared genetic aetiology between psychotic experiences and schizophrenia, but also major depressive disorder, bipolar disorder and neurodevelopmental disorders.

## Introduction

Psychotic experiences, such as hallucinations and delusions, are features of psychiatric disorders, for example schizophrenia and bipolar disorder, but they are also reported by approximately 5%-10% of the general population^1,2^. Psychotic experiences are only considered to be symptoms of psychiatric illness if they co-occur with other features of that disorder, including some aspect of psychosocial impairment. It is currently unclear whether psychotic experiences in the general population are (i) on a spectrum that at the extreme relates specifically to schizophrenia, (ii) largely unrelated to the psychotic symptoms experienced in schizophrenia and other major mental disorders, or (iii) related to liability to major mental disorders more generally.

Twin studies and genome-wide association studies (GWAS) have provided evidence that psychotic experiences are heritable (30-50% from twin studies^3-5^, 3-17% for SNP-heritability estimates^6,7^), indicating that common genetic variants play a role in their aetiology. There have been three GWAS of psychotic experiences to date, all conducted in adolescent samples^7-9^ with relatively small sample sizes (largest total sample *n*=10,098) and no reported genome-wide significant findings. These studies have focused on adolescence under the hypothesis that psychotic experiences at this age could be associated with an increased risk of mental health disorders in later life^10^. Whilst there was an initial assumption that psychotic experiences in adolescence would specifically increase the risk for schizophrenia, evidence suggests a non-specific increased risk for a broader psychopathology^10^, suggesting that research on psychotic experiences may have an important role in understanding the pathway to a wide array of clinical diagnoses^5^. However, to date no study has found strong evidence for association between genetic liabilities for schizophrenia or any other mental disorder with psychotic experiences^7,8,11-13^.

Although many individuals with a lifetime history of psychotic experiences have their first experience in adolescence, nearly a quarter of first onset psychotic experiences occur after 40 years of age^14^. It is possible that psychotic experiences first occurring in adulthood have a different aetiology to those occurring in adolescence. Our aims were to use the UK Biobank to (i) identify genetic loci associated with psychotic experiences reported by adults in a population-based study, and (ii) to determine whether genetic liability to psychotic experiences is shared with schizophrenia and/or other neuropsychiatric disorders and traits.

## Methods

### Sample

Study individuals were from the UK Biobank, a large prospective population-based cohort study of approximately 500,000 individuals aged between 40-69 who were recruited from 22 assessment centres across the UK between 2006-2010^15^. The North West Multi-Centre Ethics Committee granted ethical approval to UK Biobank and all participants provided informed consent. This study was conducted under UK Biobank project numbers 13310 and 14421.

### Psychotic experiences phenotypes

A Mental Health Questionnaire (MHQ) was sent to all participants in 2016-2017 who provided an email address and was completed by a total of 157,397 individuals (46% of those emailed, 31% of the total UK Biobank sample). For psychotic experiences, participants were asked about previous experience of visual hallucinations, auditory hallucinations, delusions of reference, delusions of persecution, how often these experiences occurred and how distressing they found them. The questions asked are detailed in **Supplementary Methods**. All analyses were conducted excluding individuals with a diagnosis of a psychotic disorder (schizophrenia, bipolar disorder or any other psychotic disorder) as defined in **Supplementary Methods.**

We selected three primary phenotypes for GWAS: (i) *any psychotic experience* defined as a positive response to any of the four symptom questions (UKB field IDs: 20471, 20463, 20474, 20468), (ii) a *distressing psychotic experience,* defined as *any psychotic experience* that was rated as ‘a bit’, ‘quite’ or ‘very’ distressing (UKB field ID: 20462), (iii) *multiple occurrence psychotic experiences* defined as *any psychotic experience* that occurred on more than one occasion (UKB field IDs: 20473, 20465, 20476, 20470). As a comparator group, we included individuals who provided a negative response to all four psychotic experience symptom questions. **Supplementary Figure 1** details the overlap in samples between these phenotypes.

We investigated additional psychotic experience phenotypes to those described above for association with PRS: (i) *visual hallucination* (UKB field ID: 20471), (ii) *auditory hallucination* (UKB field ID: 20463), (iii) *delusion of reference* (UKB field ID: 20474), (iv) *delusion of persecution* (UKB field ID: 20468), and (v) *non-distressing psychotic experiences* defined as any psychotic experience rated as not distressing (UKB field ID: 20462).

### Genetic data

Genetic data for the study participants was provided by UK Biobank and the imputation and quality control procedures are fully described elsewhere^16^. The data release contained 488,377 participants assayed on either the UK Biobank Axiom or the UK BiLEVE Axiom purpose-built arrays at the Affymetrix Research Services Laboratory. Standard quality control procedures were applied prior to imputation using Haplotype Reference Consortium (HRC)^17^ and UK10K haplotype reference^18^ panels. We applied additional quality control filters to select high quality SNPs^19^; minor allele frequency (MAF) > 0.01, imputation score (INFO) > 0.8, missingness < 0.05, Hardy-Weinberg equilibrium (HWE) < 1 × 10^−6^, and removed SNPs imputed by the UK10K haplotype reference^18^ dataset in accordance with guidance from the UK Biobank (URL: http://www.ukbiobank.ac.uk/2017/07/important-note-about-imputed-genetics-data/). One member from each related pair with a kinship coefficient > 0.15 was excluded from analyses, preferentially retaining individuals that had experienced a psychotic experience, and were otherwise removed at random.

Analyses were restricted to individuals with a self-reported British and Irish background (UKB field ID: 21000) and principal components supplied by UK Biobank^16^ (UKB field ID: 22009) were used to assess and control for population structure. European genetic ancestry was assessed using the ‘covMCD’ function in the R package ‘robustbase’^20,21^ and additional exclusions applied (described in **Supplementary Methods**).

### GWAS analysis

To identify genetic risk variants for psychotic experiences, association analysis was performed in SNPTEST (v2.5.4)^22^ using bgen v1.2 imputed dosage data^23^. Over 7.5 million SNPs were included in each GWAS. An additive logistic regression model was used including as covariates genotyping array, the top five principal components (as recommended for most GWAS approaches^24^), and any additional principal components from the first 20 that were nominally associated (*p* < 0.05) with the GWAS phenotype in a logistic regression. To obtain relatively independent index SNPs, linkage disequilibrium (LD) clumping was performed in PLINK^25^ (r^2^ < 0.1, *p* < 1×10^−4^, window size < 3MB) for each GWAS using a reference panel of 1000 randomly selected UK Biobank individuals with confirmed European ancestry. Functional annotation was conducted using FUMA^26^. LDSC^27^ was used to calculate the LD score intercept and heritability on the observed scale using the summary statistics from each psychotic experience GWAS.

#### Validation analyses in ALSPAC cohort

To assess the reproducibility of the psychotic experience GWAS, we used the summary statistics from the *any psychotic experience* GWAS to target psychotic experiences in the Avon Longitudinal Study of Parents and Children (ALSPAC) cohort^28,29^, which have been previously described^11,30^. A psychotic experience PRS was generated for each ALSPAC participant using the accepted method ^31^ and logistic regression was used to test for association between PRS and psychotic experiences reported at age 12 and 18 years. See **Supplementary Methods** for further details.

### Polygenic risk scores

To examine the relationship between psychotic experiences and genetic risk for psychiatric and personality traits from publicly available GWAS datasets (that did not include UK Biobank where possible) included; schizophrenia^32^, bipolar disorder^33^, major depressive disorder^34^, neuroticism^35^, and intelligence^36^. PRSs were generated using the method described by the PGC^31^ and detailed in **Supplementary Methods**. The intelligence GWAS excluded UK Biobank participants (n=74,214 individuals remaining, summary statistics specifically derived for this study). As it was not possible to obtain the major depressive disorder GWAS summary statistics excluding UK Biobank participants, we conducted sensitivity analyses removing individuals with a diagnosis of depression to control for overlap with the discovery dataset. The primary analysis used standardised scores generated from SNPs with a discovery sample p-value threshold of *p* ≤ 0.05, but associations at 10 other p-value thresholds were also tested. A logistic regression model was used to test the association of each PRS with various psychotic experience phenotypes, covarying for the first five principal components and genotyping array.

### Genetic correlations

LDSC^27,37^ was used to calculate the genetic correlation between each psychotic experience GWAS and psychiatric and personality traits. External GWAS datasets used to generate the correlations were the same as those used for PRS analysis (detailed above) but also included attention deficit/hyperactivity disorder (ADHD)^38^, autism spectrum disorder^39^, and the PGC cross-disorder analysis^40^. A Bonferroni correction was applied to control for multiple testing.

### Copy number variation

CNV calling, which has been described in detail elsewhere^41^, was carried out using biallelic markers common to both genotyping platforms using PennCNV-Affy^42^. Exclusion criteria included ≥ 30 CNVs, waviness factor <-0.03 or > 0.03, call rate < 0.96 for individual samples, LRR SD > 0.35 and coverage of < 20 probes, density coverage of < 1 probe per 20,000 base pairs or a confidence score of < 10 for individual CNVs. We compared carrier status of rare CNVs previously associated with (i) schizophrenia^43^ and (ii) neurodevelopmental disorders more widely^44^ (which includes all schizophrenia-associated CNVs and excludes 15q11.2 duplication because of its high frequency following our previous publications^41^) with the three primary psychotic experience phenotypes used for GWAS. Association analyses were carried out using logistic regression and included age, sex and genotyping array as covariates.

## Results

A total of 7803 (5.0%, 60.0% female) individuals from UK Biobank who completed the MHQ reported at least one psychotic experience, 3012 individuals rated the psychotic experience as distressing, and a total of 4388 reported multiple occurrences of at least one psychotic experience. A total of 147,461 individuals (56.0% female) who reported no psychotic experiences constituted the comparison group for our association analyses. The mean age of the first psychotic experience was 31.6 years (SD=17.6) (**Supplementary Figure 4**) with 2341 (35.2%) first occurring before the age of 20 or for as long as the participant could remember, 2137 (32.1%) between the ages of 20-39, and 2176 (32.7%) between the ages of 40 and 76. We excluded 198 individuals who had a diagnosis of schizophrenia, 818 with bipolar disorder, and 346 with other psychotic disorders from all genetic analyses (detailed in **Supplementary Methods)**.

### GWAS

The GWAS of *any psychotic experience* in 6123 cases and 121,843 controls (following QC, exclusions detailed in **Supplementary Methods**) identified two variants that were associated at the genome-wide significance (GWS) level of *p* < 5×10^−8^ (**Figure 1**, **Table 1**, λ_GC_=1.05, LD Score regression intercept=1.00); rs10994278, an intronic variant within Ankyrin-3 (*ANK3*) on chromosome 10 (OR=1.16, 95% confidence intervals (CI)=1.10-1.23, *p*=3.06×10^−8^), and rs549656827, an intergenic variant on chromosome 5 (OR=0.61, 95% CI=0.50-0.73, *p*=3.30×10^−8^).

A second GWAS restricting the cases to 2143 individuals with *distressing psychotic experiences* identified two GWS variants (**Table 1**, λ_GC_=1.03, LD Score intercept=1.01); rs75459873 intronic to cannabinoid receptor 2 (*CNR2*) on chromosome 1 (OR=0.66, 95% CI=0.56-0.78, *p*=3.78×10^−8^) and rs3849810, an intergenic variant on chromosome 8 (OR=1.22, 95% CI=1.13-1.31, *p*=4.55×10^−8^).

**Figure 1.**
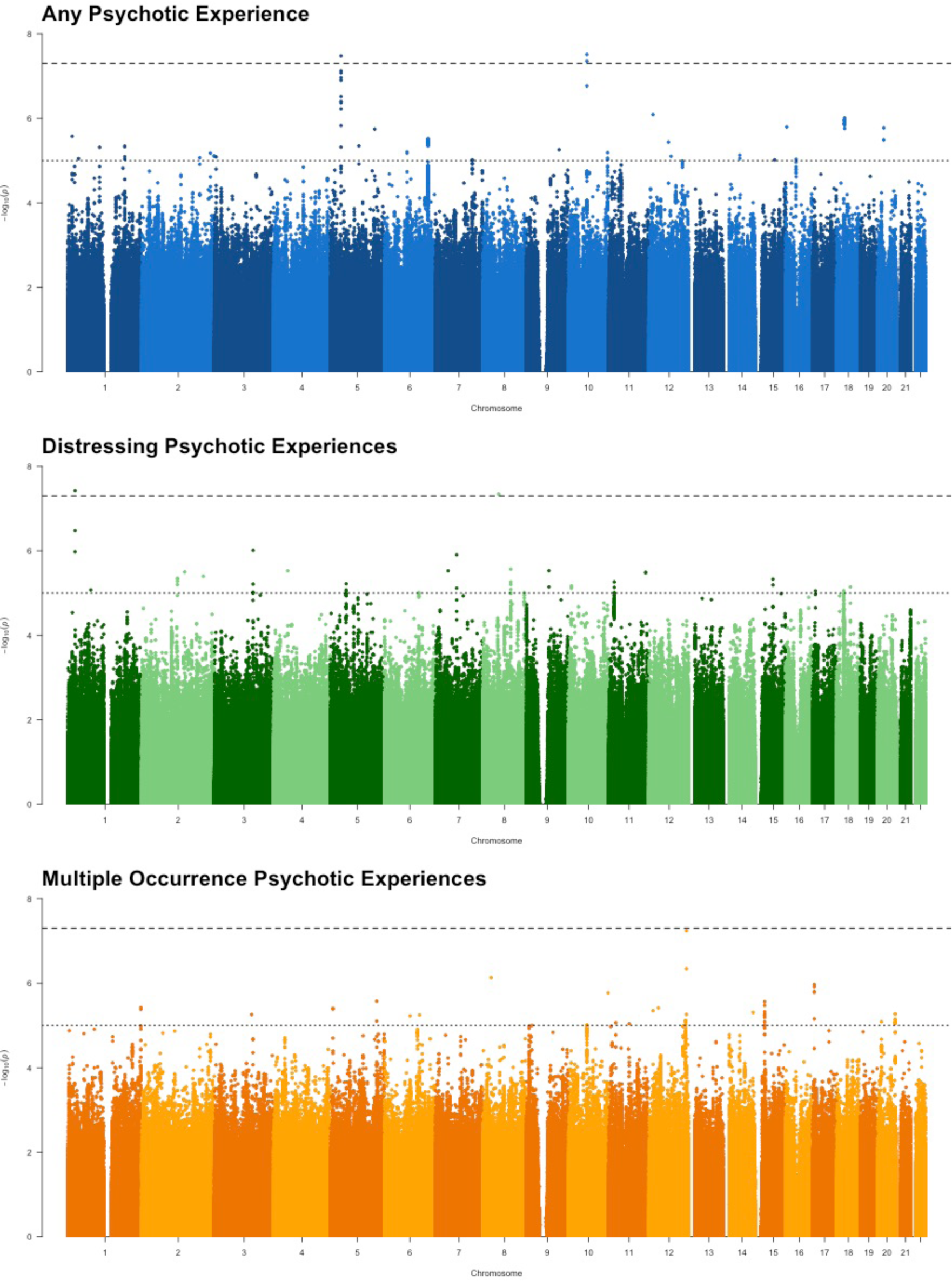
Manhattan plot for GWAS analyses of *any psychotic experience, distressing psychotic experiences* and *multiple occurrence psychotic experiences*. Dashed line represent the genome-wide significance level of *p* < 5 × 10^−8^ and the dotted line represents *p* < 1 × 10^−5^.

**Table 1.**

| Phenotype | SNP | CHR | BP | A1 | OR (95% CI) | P | Position | Nearest gene |
| --- | --- | --- | --- | --- | --- | --- | --- | --- |
| Any PE | rs10994278 | 10 | 62009219 | T | 1.16 (1.10-1.23) | $3.06 \times 10^{-8}$ | Intronic | *ANK3* |
| Any PE | rs549656827 | 5 | 35349428 | G | 0.61 (0.50-0.73) | $3.30 \times 10^{-8}$ | Intergenic | *U3, PRLR* |
| Distressing PE | rs75459873 | 1 | 24256312 | G | 0.66 (0.56-0.78) | $3.78 \times 10^{-8}$ | Intronic | *CNR2* |
| Distressing PE | rs3849810 | 8 | 53009606 | A | 1.22 (1.13-1.31) | $4.55 \times 10^{-8}$ | Intergenic | *ST18* |

Genome-wide significant associations ( $p < 5 \times 10^{-8}$ ) from GWAS analyses. Columns represent GWAS phenotype (PE = psychotic experience), variant (SNP), chromosome (CHR), base position (BP), risk allele (A1), odds ratio (OR) and 95% confidence intervals, p-value (P), position of risk variant (Position), and the nearest gene.

The third GWAS restricting the cases to 3337 individuals who reported *multiple occurrences of psychotic experiences* did not identify any associated variants at GWS (λ_GC_=1.03, LD Score intercept=1.00).

Summary statistics for SNPs associated at *p*<1×10^−5^ for each GWAS are given in **Supplementary Tables 1-3** and QQ plots are displayed in **Supplementary Figure 5**. SNP based heritability estimates calculated by LDSC^27^ for the GWAS of *any psychotic experience* was h^2^=1.71% (95% CI=1.02-2.40%). The other GWAS analyses did not requirements (case *n*>5,000 *and* Z-score>4^45^) for heritability or genetic correlation analyses with LDSC.

#### Validation analyses in ALSPAC

There was evidence of association between the PRS calculated using the *any psychotic experience* GWAS from UK Biobank at the p-value threshold of ≤0.5 and definite psychotic experiences between ages 12 and 18 years in ALSPAC (OR=1.13; 95% CI=1.02-1.25; R^2^=0.002; *p*=0.02). This finding was consistent for thresholds below *p*<0.05 and when using measures from age 18 years only (**Supplementary Figure 6**, all results detailed in **Supplementary Table 4**). However, the psychotic experiences PRS was also associated with the presence of major depressive disorder at age 18 (OR = 1.19; 95% CI=1.05-1.35; R^2^=0.004; *p*=0.01).

#### Relationship between association at *CNR2* and cannabis use

We investigated the possibility of either a mediating or moderating effect of cannabis use on the association between *distressing psychotic experiences* and rs75459873 at the cannabinoid receptor gene *CNR2*. Cannabis use itself (UKB field id: 20453) was significantly associated with *distressing psychotic experiences* (OR=1.36, 95% CI=1.32-1.40, *p*=9.16 × 10^−88^), but rs75459873 was not associated with cannabis use (OR=0.99, 95% CI=0.95-1.04, *p*=0.70). Furthermore, the association between rs75459873 and *distressing psychotic experiences* (OR=0.62, 95% CI=0.52-0.75) was unchanged in a model including cannabis use as a covariate (OR=0.62, 95% CI=0.52-0.75) or as an interaction term (OR=0.59, 95% CI=0.48-0.73, interaction *p*=0.31). Thus, we found no indication of either a mediating or moderating effect of cannabis use on the association between rs75459873 and *distressing psychotic experiences*.

### Polygenic risk score analysis

We found evidence of a weak association between *any psychotic experience* and genetic liability indicated by PRS at a *p*<0.05 threshold for SNP inclusion for schizophrenia (OR=1.09; 95% CI=1.06-1.12; adjusted R^2^=0.001; *p*=2.96×10^−11^), major depressive disorder (OR=1.16; 95% CI=1.13-1.19; R^2^=0.003; *p*=1.48×10^−30^), and bipolar disorder (OR=1.07; 95% CI=1.04-1.10; R^2^=0.001; *p*=5.11×10^−7^). These findings were consistent across most p-value thresholds (**Figure 2** and **Supplementary Figure 7**). **Supplementary Figure 8** compares the average PRS scores (SNP inclusion threshold *p*<0.05) for each psychotic experience phenotype. Individuals with *distressing psychotic experiences* had significantly higher PRS scores for schizophrenia, bipolar disorder and major depressive disorder than those with *non-distressing psychotic experiences* (**Supplementary Table 6**).

**Figure 2.**
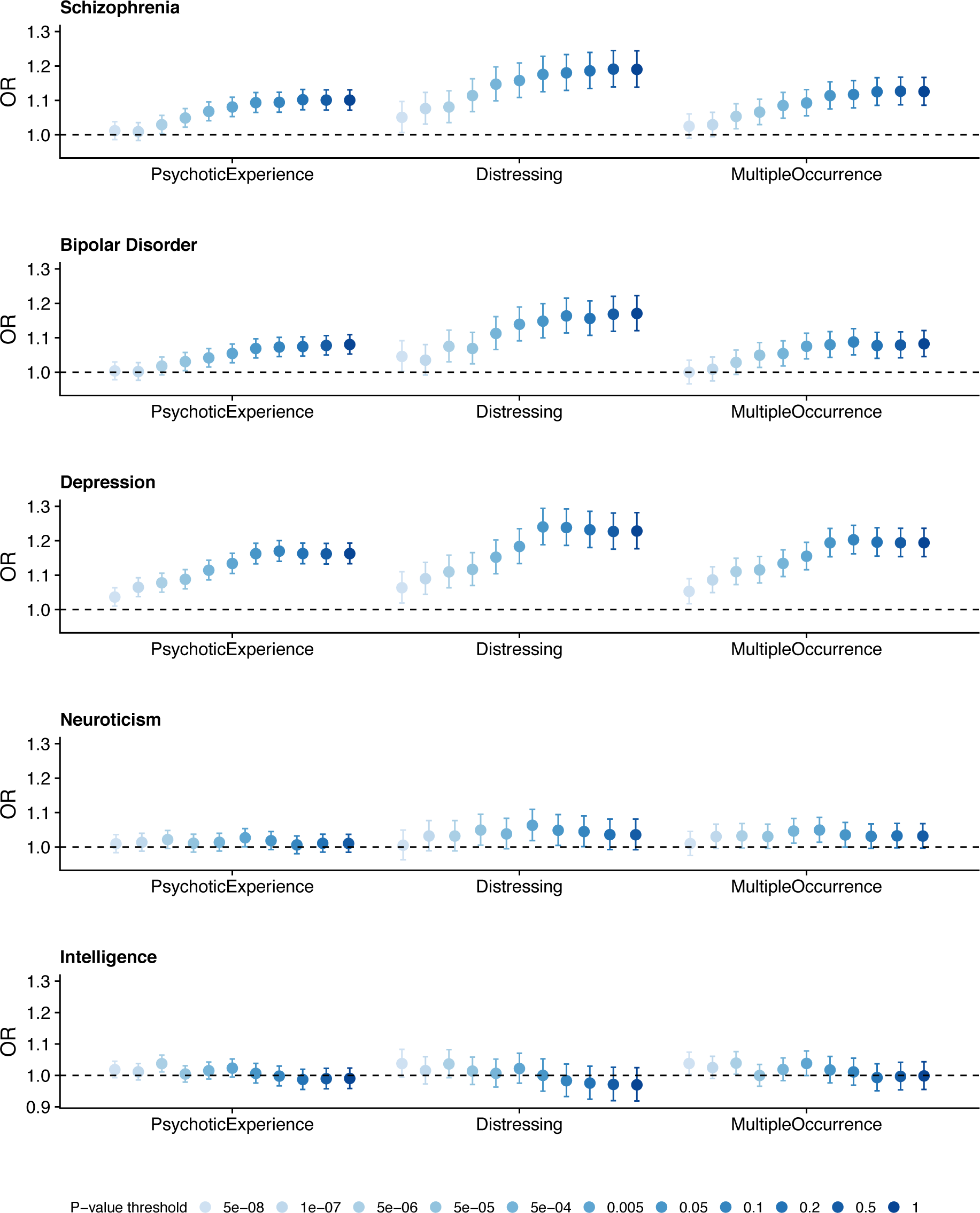
Polygenic risk score analysis. Points represent odds ratio (OR) and error bars are 95% confidence intervals. A plot with r^2^ and p-values is presented in *Supplementary Figure 7.*

We also considered individual psychotic symptoms and found that PRS for schizophrenia was more strongly associated with delusions of persecution than the other psychotic symptoms measured (**Supplementary Table 6**). This pattern was similar for PRS related to bipolar disorder and major depressive disorder and was consistent across p-value thresholds (**Supplementary Figure 9-10**). The association with psychotic experience phenotypes for bipolar disorder PRS and major depressive disorder PRS remained significant in analyses controlling for schizophrenia PRS (at *p*<0.05 threshold, **Supplementary Table 7**). The association with major depressive disorder PRS remained significant when individuals with a diagnosis of depression were removed to control for any potential overlap with the discovery dataset (**Supplementary Table 8**).

### Genetic correlations

Significant genetic correlations (r_g_) were observed between *any psychotic experience* and major depressive disorder (r_g_=0.46, *p*=4.64×10^−11^), autism spectrum disorder (r_g_=0.39, *p*=1.68×10^−4^), the PGC cross-disorder GWAS (r_g_=0.30, *p*=1.53×10^−3^) and schizophrenia (r_g_=0.21, *p*=7.29×10^−5^). **Figure 3** displays these genetic correlations and **Supplementary Table 5** details the full results. The other GWAS analyses did not meet requirements^45^ for genetic correlation analysis.

**Figure 3.**
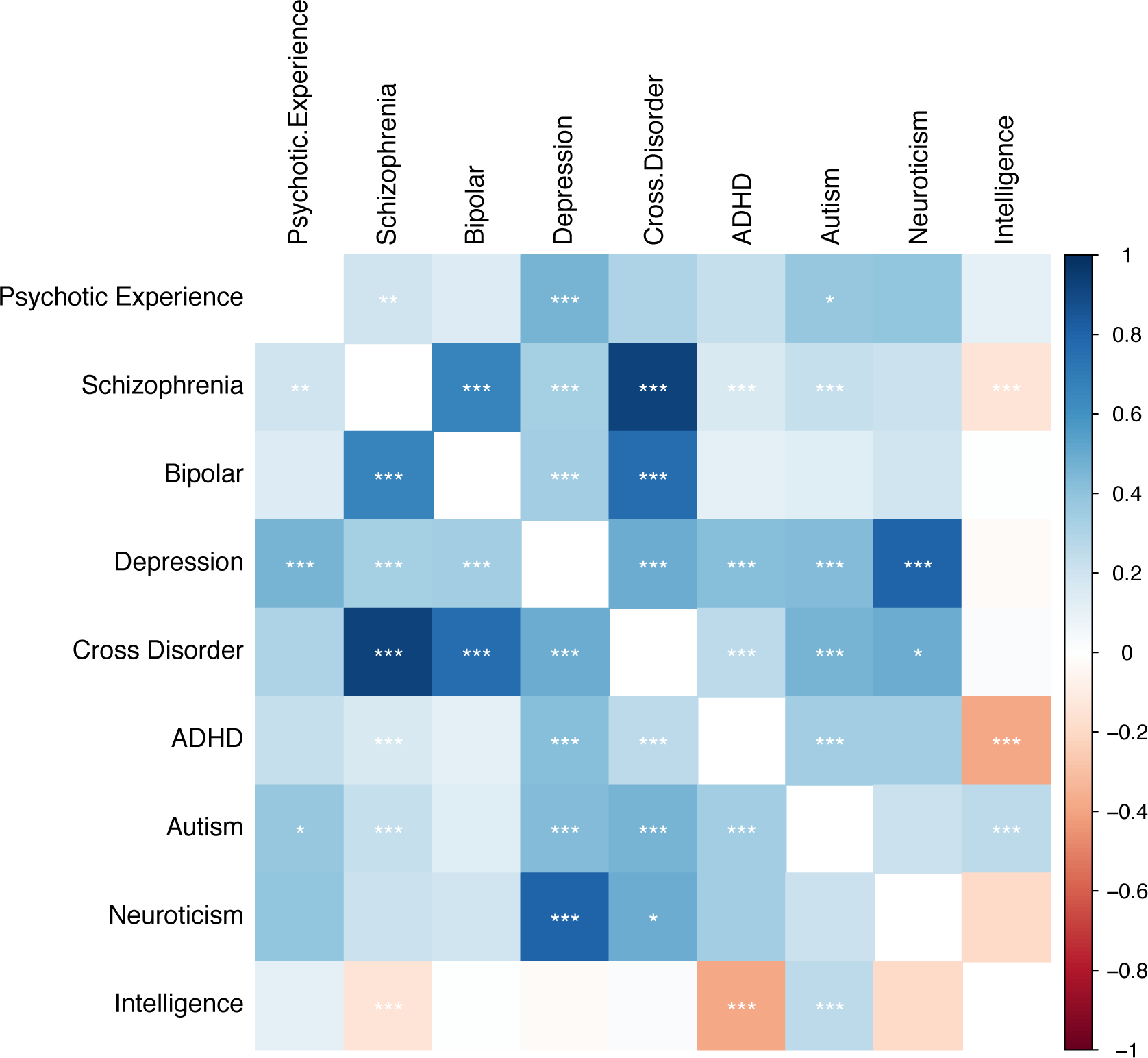
Genetic correlation analysis. Colour corresponds to the strength of the correlation (rg) and stars correspond to the statistical significance of the correlation (* P < 0.003, ** P < 0.0001, *** P < 0.00001). Positive correlations are shown in blue and negative correlations in red.

### Copy number variation

Individuals reporting *distressing psychotic experiences* in particular had an increased burden of CNVs previously associated with schizophrenia (OR=2.04; 95% CI=1.39-2.98; *p*=2.49×10^−4^) and neurodevelopmental disorders (OR=1.75; 95% CI=1.24-2.48; *p*=1.41×10^−3^). There was evidence of an increased burden of these CNVs in individuals reporting *any psychotic experience* but not *multiple occurrence psychotic experiences* (**Table 2**).

**Table 2.**

| *Schizophrenia CNVs* |  |  |  |  |
| --- | --- | --- | --- | --- |
|  | **Case rate (n)** | **Control rate (n)** | **OR (95% CI)** | **P** |
| Any PE | 0.010 (59/5829) | 0.006 (572/115807) | 1.54 (1.18-2.01) | $1.60 \times 10^{-3}$ |
| Distressing PE | 0.014 (28/2046) | 0.006 (572/115807) | 2.04 (1.39-2.98) | $2.49 \times 10^{-4}$ |
| Multiple occurrence PE | 0.009 (28/3177) | 0.006 (572/115807) | 1.34 (0.91-1.94) | 0.144 |
| *Neurodevelopmental disorder CNVs* |  |  |  |  |
|  | **Case rate (n)** | **Control rate (n)** | **OR (95% CI)** | **P** |
| Any PE | 0.014 (83/5829) | 0.009 (1058/115807) | 1.54 (1.23-1.92) | $1.89 \times 10^{-4}$ |
| Distressing PE | 0.017 (34/2046) | 0.009 (1058/115807) | 1.75 (1.24-2.48) | $1.41 \times 10^{-3}$ |
| Multiple occurrence PE | 0.012 (38/3177) | 0.009 (1058/115807) | 1.30 (0.92-1.78) | 0.139 |

Association of psychotic experience phenotypes with copy number variants (CNVs) previously associated with (i) schizophrenia and (ii) neurodevelopmental disorders (note all schizophrenia-associated CNVs are also included in neurodevelopmental disorders). Columns represent; phenotype (PE = psychotic experience), rate of CNVs in cases (those reporting a psychotic experience), rate of CNVs in controls (the comparator group, those who did not report a psychotic experience), odds ratio (OR) and 95% confidence intervals (CI), and p-value (P).

## Discussion

We conducted the largest GWAS of psychotic experiences using the population-based UK Biobank sample, and identified four genome-wide significant loci; an intronic variant to *ANK3*, an intronic variant to *CNR2,* and two intergenic loci. PRS, genetic correlation and CNV analyses indicated a shared genetic aetiology between psychotic experiences and schizophrenia, but also neurodevelopmental disorders and other psychiatric disorders such as major depressive disorder and bipolar disorder. The shared genetic aetiology between psychotic experiences and major mental health conditions was stronger for psychotic experiences rated as distressing.

The primary GWAS findings are related to intronic variants in *ANK3* and *CNR2*. The GWAS of *any psychotic experience* identified two significant loci, the most significant of which was indexed by rs10994278, an intronic variant to *ANK3* (OR=1.16, *p*=3.06×10^−8^). The *ANK3* gene encodes ankyrin-G, a protein that has been shown to regulate the assembly of voltage-gated sodium channels and is essential for normal synaptic function^46^. *ANK3* is one of strongest and most replicated genes for bipolar disorder^33^, and variants within *ANK3* have also been associated in the PGC cross-disorder GWAS^40^, and in a rare variant analysis of autism spectrum disorder^47^.

The GWAS of *distressing psychotic experiences* also identified two significant loci, the most significant of which was indexed by rs75459873, an intronic variant to *CNR2* (OR=0.66, *p*=3.78×10^−8^). *CNR2* encodes for CB2, one of two well-characterised cannabinoid receptors (CB1 being the other). CB2 receptors they have abundant expression in the immune system, microglia and some neurons^48^. Several lines of evidence have implicated the endocannabinoid system in psychiatric disorders including schizophrenia^49,50^ and depression^51^. The main psychoactive agent of cannabis, Δ^9^-tetrahydrocannabinol, can cause acute psychotic symptoms and cognitive impairment^52^. Given that cannabis use is strongly associated with psychotic experiences, we tested, but found no evidence for, a mediating or moderating effect of cannabis use on the association of rs75459873 and *distressing psychotic experiences.* However, whilst no evidence was found in this study, a mediating effect of cannabis use cannot be ruled out given there may be questionable reliability in self-reported illicit cannabis use.

We found that psychotic experiences shared genetic risk with several psychiatric disorders, which included, but was not specific to, schizophrenia. PRS analyses identified associations between psychotic experiences and genetic liability for schizophrenia, major depressive disorder and bipolar disorder but not for intelligence or neuroticism. The magnitude of associations in the PRS analysis was comparable between psychiatric disorders and was very small (maximum r^2^ value<1% and AUC<0.55). We found particular enrichment of schizophrenia, major depressive disorder and bipolar disorder PRS in cases that found the psychotic experiences distressing and for delusions of persecution. Genetic correlation analysis identified significant genetic correlations between psychotic experiences and major depressive disorder (r_g_=0.46), schizophrenia (r_g_=0.21), autism spectrum disorder (r_g_=0.39), and for the PGC cross-disorder GWAS (r_g_=0.30). Furthermore, we found an increased burden of CNVs previously associated with schizophrenia (OR=2.04) and neurodevelopmental disorders more widely (OR=1.75) in individuals with distressing psychotic symptoms. There was also an increased burden of CNVs in those reporting any psychotic experience but the association was stronger for distressing psychotic experiences. All schizophrenia-associated CNVs are also associated with neurodevelopmental disorders such as intellectual disability and autism spectrum disorder, and in fact penetrance is higher in these disorders^43^. Furthermore, CNVs in UK Biobank have been associated with a range of outcomes including cognitive performance^41,53^ and depression^54^, adding strength to our findings of a lack of specificity for psychotic experiences genetic risk.

A number of studies have demonstrated that psychopathology in the population is best described by a bifactor model with a common latent trait as well as specific traits, and that psychotic experiences index the more severe end of the common or shared trait^56,57^. Our findings of non-specificity of genetic risk for psychotic experiences with risk for other disorders are consistent with previous studies^7,8,11,12^. Nonetheless, despite lacking specificity, our results suggest that incorporating questions about frequency and, in particular, distress of psychotic experiences to self-reported assessments may allow a more valid identification of experiences that index schizophrenia and major mental health disorder liability.

Given the heritability estimate was very low (1.7%) for the *any psychotic experience* GWAS and the low amount of variance explained in our PRS analysis, and those of others^7,11^, our findings indicate that understanding the genetics of psychotic experiences is unlikely to have an important impact on understanding the genetics of schizophrenia specifically. However, given that the small heritability of psychotic experiences is substantially shared with other psychiatric disorders, and that non-genetic aetiology of psychotic experiences has been shown to be shared^55^; then studies of psychotic experience aetiology may be informative for psychiatric disorders in general.

### Strengths and limitations

Strengths of this study include the large sample size (approximately 10x that of previous studies), which is required for genetic association studies, the use of an adult cohort, and the use of multiple psychotic experience phenotypes, which increase confidence in our findings. Furthermore, individuals with psychotic disorders were removed from our analyses, and thus the results will not be driven by this smaller subset of individuals.

One limitation of this study relates to the retrospective measurement of lifetime psychotic experiences by self-report from an online questionnaire, as this increases the likelihood of measurement error. A further limitation is the evidence of a ‘healthy volunteer’ bias for the participants recruited to UK Biobank and the sample cannot be therefore considered representative of the general population^58^. We also found that the participants that completed the MHQ had significantly higher intelligence and lower schizophrenia, depression and neuroticism PRS compared to UK Biobank participants who did not complete the MHQ (**Supplementary Table 9**). It is possible therefore that our results are affected by selection bias. Lastly, we were not able to entirely de-duplicate the UK Biobank individuals from all of the external datasets used for the polygenic risk score analysis. We excluded datasets that explicitly included the UK Biobank where possible, and failing that removed cases in sensitivity analyses.

### Conclusions

In the largest GWAS of psychotic experiences from the population-based UK Biobank sample, we identified four genome-wide significant loci associated with psychotic experiences including an intronic variant to *ANK3*, an intronic variant to *CNR2,* and two intergenic loci. We found support for a shared genetic aetiology between psychotic experiences and several psychiatric disorders including schizophrenia, major depressive disorder, bipolar disorder, and neurodevelopmental disorders indicating that psychotic experiences are not specifically related to schizophrenia, but rather to a general risk for mental health disorders.

## Acknowledgements

The National Institute for Health Research, Medical Research Council and British Heart Foundation supplied funds to complete genotyping on all UK Biobank participants. The funding bodies had no experimental role in the comparison between DNA quantification methods used in the genotyping workflow. We thank UK Biobank participants for providing the samples for this project.

This project was supported by the following grants: Medical Research Council (MRC) Centre (MR/L010305/1), Program (G0800509), Project (MR/L011794/1) grants to Cardiff University, and The National Centre for Mental Health (NCMH), funded by the Welsh Government through Health and Care Research Wales. HJJ and SZ are supported by the NIHR Biomedical Research Centre at University Hospitals Bristol NHS Foundation Trust and the University of Bristol. KK is supported by a Wellcome Trust Clinical Research Training Fellowship. KD and MH are supported by the National Institute for Health Research (NIHR) Biomedical Research Centre at South London and Maudsley NHS Foundation Trust and King’s College London. The views expressed are those of the author(s) and not necessarily those of the NHS, the NIHR or the Department of Health and Social Care.

The UK Medical Research Council, the Wellcome Trust (Grant ref: 102215/2/13/2) and the University of Bristol provide core support for ALSPAC. A comprehensive list of grants funding is available on the ALSPAC website. The collection of ALSPAC measures used was specifically funded by the Medical Research Council (Grant ref: G0701503/85179). ALSPAC GWAS data was generated by Sample Logistics and Genotyping Facilities at the Wellcome Trust Sanger Institute and LabCorp (Laboratory Corporation of America) using support from 23andMe. We are extremely grateful to all the ALSPAC families who took part in this study, the midwives for their help in recruiting them, and the whole ALSPAC team, which includes interviewers, computer and laboratory technicians, clerical workers, research scientists, volunteers, managers, receptionists and nurses.

