## Supplementary Methods for "Genetic association study of psychotic experiences in UK Biobank"

### *Supplementary Information*

Genetic association study of psychotic experiences in UK Biobank	1

Supplementary Information	1

Supplementary Methods	2

Unusual and psychotic experience questions	2

Defining European genetic ancestry	4

Defining schizophrenia, bipolar disorder and psychotic disorder	4

Calculation of polygenic risk scores in UK Biobank	4

Replication of psychotic experiences GWAS in ALSPAC cohort	5

Details of individuals excluded	8

Supplementary Figure 1: Phenotypes	10

Supplementary Figure 2: PCA analysis	11

Supplementary Figure 3: PCA analysis	12

Supplementary Figure 4: Age of onset of PEs	13

Supplementary Figure 5: QQ plots of GWAS and gene analysis	14

Supplementary Figure 6: PRS analysis of PE symptoms in ALSPAC	15

Supplementary Figure 7: PRS analysis of PE phenotypes	16

Supplementary Figure 8: Average polygenic risk by PE phenotype	17

Supplementary Figure 9: PRS analysis of PE symptoms	18

Supplementary Figure 10: PRS analysis of PE symptoms	19

References	20

### Supplementary Methods

#### Unusual and psychotic experience questions

The following questions are from Section F of the Mental Health Questionnaire (MHQ) administered via web-based questionnaire for UK Biobank participants.

| **Q. No** | **Question** | **Responses** |
| --- | --- | --- |
| INTRO | The next set of questions is about unusual experiences that you may have had, like seeing visions or hearing voices. We believe that these things may be quite common, but we don't know for sure. So please take your time and think carefully before answering. |  |
| F1 | Did you ever see something that wasn’t really there that other people could not see?  Please do not include any times when you were dreaming or half-asleep or under the influence of alcohol or drugs. | [Choose one from]  - 01 Yes  - 00 No  - NA Do not know  - DA Prefer not to answer |
| F1a | About how many times in your life did this happen (when you were not dreaming, not half-asleep, and not under the influence of alcohol or drugs)? | FBOX1: Integer box 1 – 999  FBOX1 & “time(s)”  OR  - 01 Too many to count  - NA Do not know  - DA Prefer not to answer |
| F2 | Did you ever hear things that other people said did not exist, like strange voices coming from inside your head talking to you or about you, or voices coming out of the air when there was no one around?  Please do not include any times when you were dreaming or half-asleep or under the influence of alcohol or drugs. | [Choose one from]  - 01 Yes  - 00 No  - DA Prefer not to say  - NA Don’t know |
| F2a | About how many times in your life did this happen (when you were not dreaming, not half-asleep, and not under the influence of alcohol or drugs)? | FBOX2: Integer box 1 – 999  FBOX2 & “time(s)”  OR  - 01 Too many to count  - NA Do not know  - DA Prefer not to answer |
| F3 | Did you ever believe that a strange force was trying to communicate directly with you by sending special signs or signals that you could understand but that no one else could understand (for example through the radio or television)?  Please do not include any times when you were dreaming or half-asleep or under the influence of alcohol or drugs. | [Choose one from]  - 01 Yes  - 00 No  - NA Do not know  - DA Prefer not to answer |
| F3a | About how many times in your life did this happen (when you were not dreaming, not half-asleep, and not under the influence of alcohol or drugs)? | FBOX3: Integer box 1 – 999  FBOX3 & “time(s)”  OR  - 01 Too many to count  - NA Do not know  - DA Prefer not to answer |
| F4 | Did you ever believe that that there was an unjust plot going on to harm you or to have people follow you, and which your family and friends did not believe existed?  Please do not include any times when you were dreaming or half-asleep or under the influence of alcohol or drugs. | [Choose one from]  - 01 Yes  - 00 No  - NA Do not know  - DA Prefer not to answer |
| F4a | About how many times in your life did this happen (when you were not dreaming, not half-asleep, and not under the influence of alcohol or drugs)? | FBOX4: Integer box 1 – 999  FBOX4 & “time(s)”  OR  - 01 Too many to count  - NA Do not know  - DA Prefer not to answer |
| F5 | How often did any of these experiences happen in the past 1 year (seeing a vision, hearing a voice, or believing that something strange was trying to communicate with you, or there was a plot against you)? | [Choose one from]  - 00 Not at all  - 01 Once or twice  - 02 Less than once a month  - 03 More than once a month  - 04 Nearly every day or daily  - DA Prefer not to answer |
| F6 | How old were you (approximately) when you first had one of these experiences (seeing a vision, hearing a voice, or believing that something strange was trying to communicate with you, or there was a plot against you)? | FBOX5: Integer box 2 to  current age  FBOX5 & “years old”  OR  - 01 As long as I can remember  - NA Do not know  - DA Prefer not to answer |
| F7 | How distressing did you find having any of these experiences (seeing a vision, hearing a voice, or believing that something strange was trying to communicate with you, or there was a plot against you)? | [Choose one from]  - 00 Not distressing at all, it was a positive experience  - 01 Not distressing, a neutral experience  - 02 A bit distressing  - 03 Quite distressing  - 04 Very distressing  - NA Do not know  - DA Prefer not to answer |
| F8 | Did you ever talk to a doctor, counselor, psychiatrist or other health professional about any of these experiences (seeing a vision, hearing a voice, or believing that something strange was trying to communicate with you, or there was a plot against you)? | [Choose one from]  - 01 Yes  - 00 No  - NA Do not know  - DA Prefer not to answer |
| F9 | Were you ever prescribed a medication by a health professional for any of these experiences (seeing a vision, hearing a voice, or believing that something strange was trying to communicate with you, or there was a plot against you)? | [Choose one from]  - 01 Yes  - 00 No  - NA Do not know  - DA Prefer not to answer |

#### Defining European genetic ancestry

Analyses were restricted to individuals with a self-reported British and Irish ethnic background (UKBB field ID: 21000). The first 40 principal components supplied by UK Biobank^1^ (UKBB field ID: 22009) were used to assess and control for population structure. Initial analyses (**Supplementary Figure 2**) showed that filtering by self-reported ancestry does not adequately control for ancestry, and so European genetic ancestry was confirmed using the ‘covMCD’ function in the R package ‘robustbase’^2^, which uses the first five principal components to compute a Minimum Covariance Determinant (MCD) estimator of location and scatter via the deterministic MCD algorithm^3^. The MCD defines a hyper-ellipsoid in a multi-dimensional space that contains the majority of points representing individuals in that group, and once this has been defined, all individuals are allocated a distance to the hyper-ellipsoid in PC space^4^. We selected individuals that were within the 90^th^ percentile of MCD distance (**Supplementary Figure 3**).

#### Defining schizophrenia, bipolar disorder and psychotic disorder

We searched for evidence of a diagnosis of schizophrenia, bipolar affective disorder, and psychotic disorder from numerous sources within UK Biobank. Individuals were classed as having one of these disorders if there was any indication from any of the following sources (i) self-reported diagnosis at the assessment centre interview (UKBB field ID: 20002), (ii) an ICD-10 primary (UKBB field ID: 41202) or secondary (UKBB field ID: 41204) diagnosis from linked hospital records, (iii) an ICD-10 diagnosis from death records (UKBB field IDs: 40001 and 40002), or (iv) a self-report of a relevant diagnosis made by a health professional in the mental health questionnaire (MHQ) (UKBB field ID: 20544). The ICD-10 codes used for schizophrenia consisted of F20 and F25, for bipolar affective disorder F30 and F31, and for psychotic disorder F21, F22, F23, F28 and F29.

#### Calculation of polygenic risk scores in UK Biobank

To examine the relationship between PEs and genetic risk for schizophrenia, bipolar disorder, depression, neuroticism and intelligence, PRSs were generated using the method described by the PGC^31^. External discovery datasets used to generate the PRSs included the largest meta-analysis of schizophrenia to date^35^, the PGC2 GWAS of bipolar disorder^36^, latest PGC GWAS of major depressive disorder^37^, Genetics of Personality Consortium (GPC2) GWAS of neuroticism^38^, and lastly for intelligence, summary stats were obtained for the latest GWAS^39^ excluding UK Biobank participants (n=74,214 individuals remaining, summary statistics specifically derived for this study). PRSs were calculated using imputation dosage data for each UK Biobank participant that passed GWAS QC measures using PRSice (v2)^40^. High quality SNPs were selected by applying filters for INFO > 0.9, MAF > 0.1, removing indels, and excluding the extended MHC region (25 MB – 35 MB). A reference panel of 1000 randomly selected UK Biobank participants was used to obtain relatively independent SNPs (r^2^ < 0.2, window size < 500kb). The first five principal components, any additional principal components from the first 20 that were nominally associated (P < 0.05) with the GWAS phenotype in a logistic regression, and genotyping array, were added as covariates for each PRS generated. Scores were generated at 11 SNP p-value thresholds (5 x 10^-8^, 1 x 10^-7^, 5 x 10^-6^, 5 x 10^-5^, 5 x 10^-4^, 0.005, 0.05, 0.10, 0.20, 0.50, 1).

Our primary analysis of PRS used standardised scores generated from SNPs with a GWAS discovery sample p-value threshold of P ≤ 0.05. Statistical association analyses were conducted in R and used a regression model to test the association of each PRS with the various PE phenotypes, and covarying for the first five principal components and genotyping array. The Nagelkerke R^2^ and area under the curve (AUC) values were adjusted for covariates. All analyses were repeated excluding those with schizophrenia, bipolar disorder or a psychotic disorder.

#### Replication of psychotic experiences GWAS in ALSPAC cohort

To assess the validity of the PE phenotype as a reproducible and biological trait, we targeted PEs in the Avon Longitudinal Study of Parents and Children (ALSPAC) longitudinal birth cohort.

##### ALSPAC study population

The initial cohort consisted of 14,541 pregnant women residing in the former Avon Health Authority area with an expected delivery date between April 1991 and December 1992^5,6^. Of these initial pregnancies, there was a total of 14,676 fetuses, resulting in 14,062 live births and 13,988 children who were alive at 1 year of age. When the oldest children were approximately 7 years of age, an attempt was made to bolster the initial sample with eligible cases who had failed to join the study originally. The total sample size for analyses using any data collected after the age of 7 is therefore 15,247 pregnancies, resulting in 15,458 fetuses. Of this total sample of 15,656 fetuses, 14,973 were live births and 14,899 were alive at 1 year of age. Please note that the study website contains details of all the data that is available through a fully searchable data dictionary and variable search tool (<http://www.bristol.ac.uk/alspac/researchers/our-data/>). Ethical approval for the study was obtained from the ALSPAC Ethics and Law Committee and the Local Research Ethics Committees. Informed consent for the use of data collected via questionnaires and clinics was obtained from participants following the recommendations of the ALSPAC Ethics and Law Committee at the time and consent for biological samples has been collected in accordance with the Human Tissue Act (2004).

##### ALSPAC genetic data

Genetic data were acquired using the Illumina HumanHap550 quad genome-wide single nucleotide polymorphism (SNP) genotyping platform from 9912 participants. Individuals were excluded from further analysis based on gender mismatches, minimal or excessive heterozygosity, disproportionate levels of individual missingness (>3%), evidence of cryptic relatedness (>10% of alleles identical by descent) and being of non-European ancestry (assessed by multidimensional scaling analysis including HapMap 2 individuals). SNPs with a minor allele frequency (MAF) of < 1%, Impute2 information quality metric of < 0.8, a call rate of < 95% or evidence for violations of Hardy-Weinberg equilibrium (p-value < 5 x 10^-7^) were removed. Imputation of the target data was performed using Impute V2.2.2 against the 1000 genomes reference panel (Phase 1, Version 3; all polymorphic SNPs excluding singletons), using 2 186 reference haplotypes (including non-Europeans). Following quality control assessment and imputation and restricting to 1 young person per family, genetic data was available for 7975 ALSPAC individuals.

##### Generating psychotic experiences polygenic risk scores in ALSPAC

Prior to construction of PE polygenic risk scores (PRSs), SNPs were removed from the analysis if they had a MAF < 0.01, an imputation quality < 0.8, or if there was allelic mismatch between samples (the alleles reported by the discovery study did not match the alleles present in the ALSPAC sample). Due to the high linkage disequilibrium (LD) within the extended major histocompatibility complex (MHC; chromosome 6: 25-34Mb) only a single SNP was included to represent this region within the analysis. Remaining SNPs were then further pruned for linkage disequilibrium (LD) using the PLINK (v1.90)^7^ --clump command to retain SNPs with an association p*-*value ≤ 0.5 and r2 < 0.25 within 500kb windows.

PRSs were constructed using the summary statistics from the primary GWAS of *any PE* in UK Biobank. PRSs were calculated for each ALSPAC individual using the PLINK (v1.07)^7^ --score command. Scores are calculated by summing the number of reference alleles present for each SNP (0, 1 or 2) weighted by the logarithm of its odds ratio for *any PE* and standardized prior to analyses.

Our primary analysis used a score generated using a list of SNPs meeting an *any PE* GWAS p-value threshold of ≤ 0.50. To assess the robustness of our findings, analyses were then repeated using PRSs based on SNPs which were associated with *any PE* at a range of GWAS p-value thresholds (p-value ≤ 1e^-6^ to ≤ 0.5).

##### Psychotic experiences in ALSPAC

The semi-structured Psychosis-Like Symptom Interview (PLIKSi)^8,9^, which draws on principles of standardized clinical examination developed for the Schedule for Clinical Assessment in Neuropsychiatry (SCAN), was used to assess psychotic experiences in ALSPAC at ages 12 and 18 years. The PLIKSi allows rating of 12 psychotic experiences including hallucinations (visual and auditory), delusions (spied on, persecution, thoughts read, reference, control, grandiosity, other) and experiences of thought interference (broadcasting, insertion and withdrawal). Any unspecified delusions elicited are also rated. Structured stem questions (e.g. “have you ever seen something or someone that other people could not see?”; “have you ever thought you were being followed or spied on?”; “Have you ever felt that thoughts are put into your mind that are not your own?”) are followed up by cross-questioning to allow the interviewer to make a decision as to whether experiences described meet SCAN criteria for a psychotic experience.

The interviewers were psychology graduates trained in assessment using the SCAN psychosis section and using the PLIKSi. Psychotic experiences were rated as not present, suspected or definitely psychotic. Unclear responses were always ‘rated down’ and symptoms were rated as definite only when a clear example was provided. At regular intervals samples of recorded interviews were also rated by a psychiatrist to ensure interviewers were rating experiences correctly. The PLIKSi shows very good inter-rater and test-retest reliability^10^.

To maximise the numbers within our sample we used data from both the interviews at ages 12 and 18 years to identify individuals were rated as having one or more definite psychotic experiences between ages 12 and 18 years, compared to no or only suspected psychotic experiences across this age range.

To assess the sensitivity of results, analyses were also performed using data from age 12 and age 18 years separately and using a different cut-off of definite or suspected psychotic experiences compared to no psychotic experiences at 12 or 18 years. A latent variable was also generated for psychotic experiences at age 16.5 years using ten items from the self-report Psychosis-Like Symptoms Questionnaire (PLIKSq)^11^. These ten items were rated on a 3-point scale (never; maybe; definitely) and assessed presence of hallucinations, delusions and thought interference since age 15. The PE latent variable was generated in Mplus (version 7.31) as previously reported^12^.

Logistic regressions were performed in Stata v15.1 to test for associations between *any PE* PRSs and ALSPAC PEs reported at age 12 and 18 years. Linear regressions were used in Mplus to test for associations between *any PE* PRSs and the latent PEs trait at age 16.5 years. No principal components were included in the PRS regression analyses, as is standard for studies in this cohort due to homogenous nature of the sample^13^.

#### Details of individuals excluded

The table below details the numbers excluded during each quality control step.

| **QC step** | **Comparison group** | **Any PE** | **Distressing PE** | **Multiple PE** |
| --- | --- | --- | --- | --- |
| 1 | 13,429 | 840 | 360 | 510 |
| 2 | 4,603 | 274 | 97 | 150 |
| 3 | 7,511 | 11 | 6 | 7 |
| 4 | 539 | 555 | 406 | 384 |
| Total remaining | 121,379 | 6,123 | 2,143 | 3,337 |

Quality control step 1 excluded individuals that did not have a self-reported White British or Irish ethnicity, those that did not have genetic data available, and those that did not pass initial genetic quality control parameters (missingness). Step 2 excluded individuals that did not have European genetic ancestry as defined by principal components (see ‘defining European genetic ancestry’ above). Step 3 excluded related individuals and finally, step 4 excluded individuals with a schizophrenia, bipolar disorder or psychotic disorder diagnosis.

### Supplementary Figure 1: Phenotypes

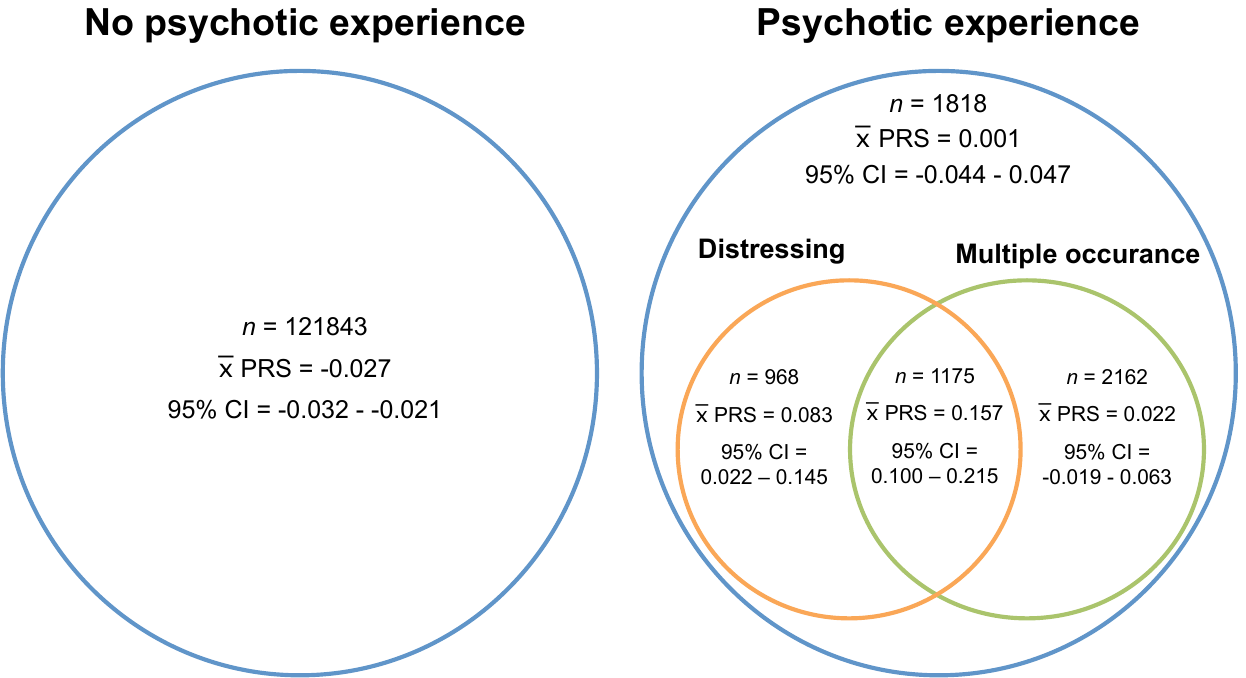

Venn diagram detailing the sample overlaps between the phenotypes used in the three psychotic experience GWAS analyses. The left hand circle represents the comparison group, who reported no psychotic experiences. The right hand circle represents the cases used in the three GWAS (i) any psychotic experience, (ii) distressing psychotic experiences, and (iii) multiple occurrence psychotic experiences. The mean polygenic risk score for schizophrenia, and 95% confidence interval for the mean polygenic risk score, is given for each subset.

### Supplementary Figure 2: PCA analysis

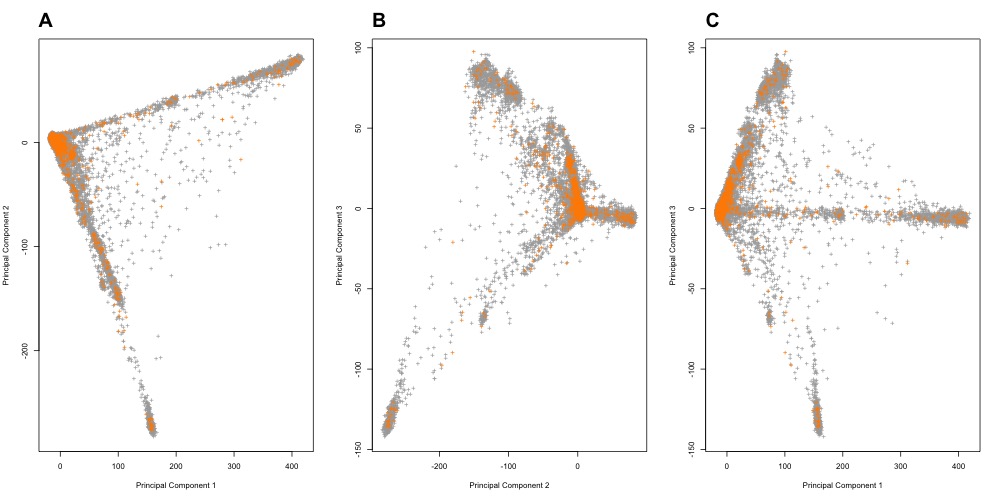

Principal component (PC) analysis. Points represent individuals who completed the MHQ, grey points: *n*=139,975 controls who completed MHQ, orange points *n*=7,791 cases with any self-reported psychotic experience. Plot A: PC 1 and PC2, plot B: PC2 and PC3, plot C: PC1 and PC2.

### Supplementary Figure 3: PCA analysis

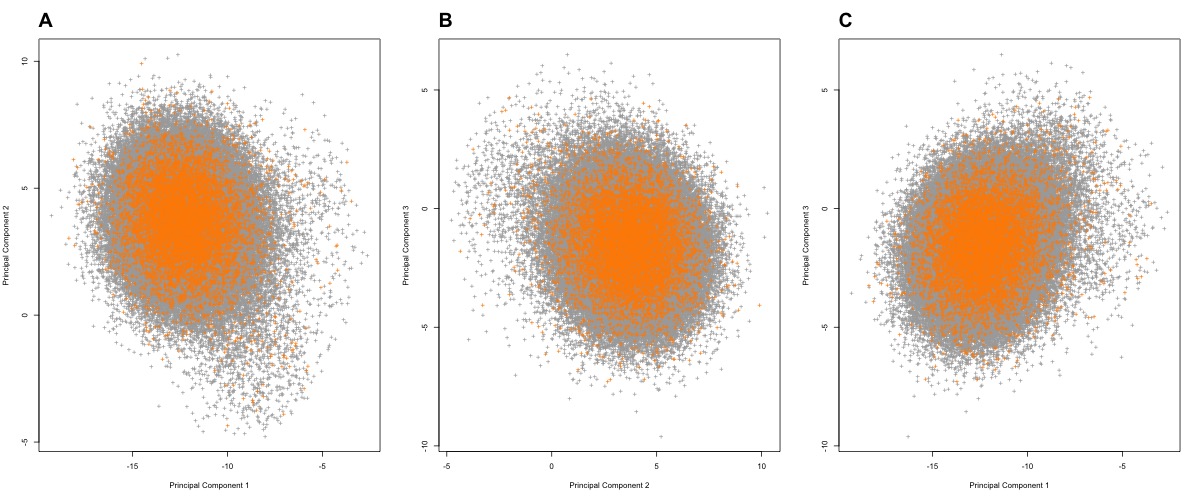

Principal component (PC) analysis after exclusions from MCD model. Points represent individuals who completed the MHQ, grey points: controls who completed MHQ, orange points: cases with any self-reported psychotic experience. Plot A: PC 1 and PC2, plot B: PC2 and PC3, plot C: PC1 and PC2.

### Supplementary Figure 4: Age of onset of PEs

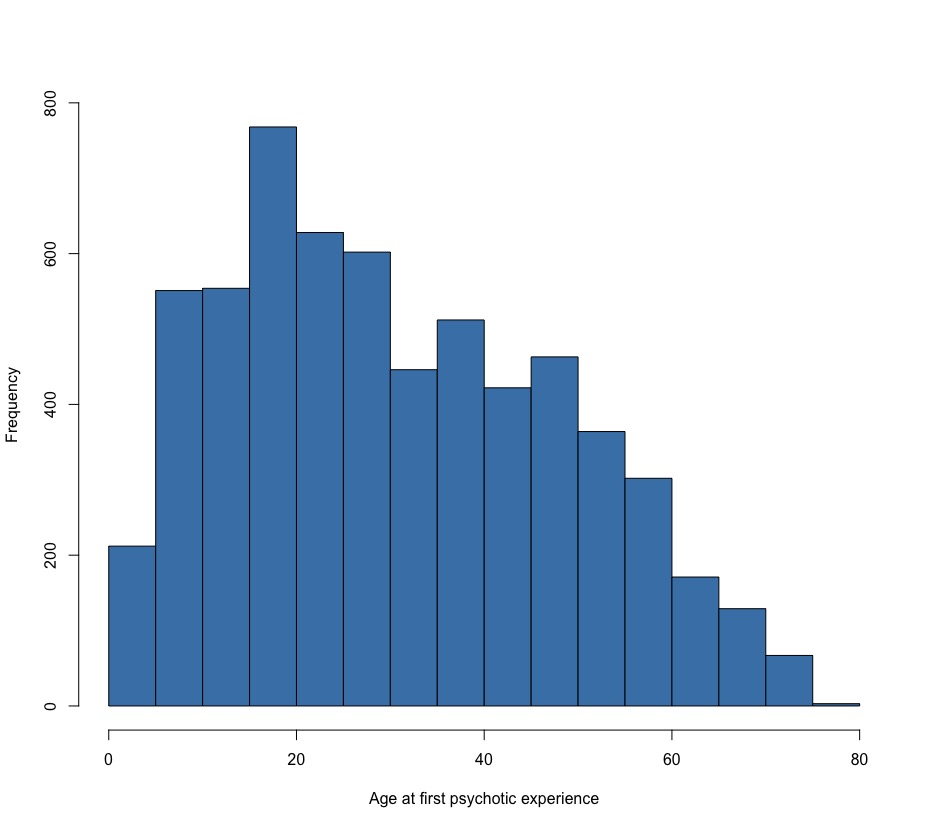

Histogram of age at first psychotic experience.

### Supplementary Figure 5: QQ plots of GWAS and gene analysis

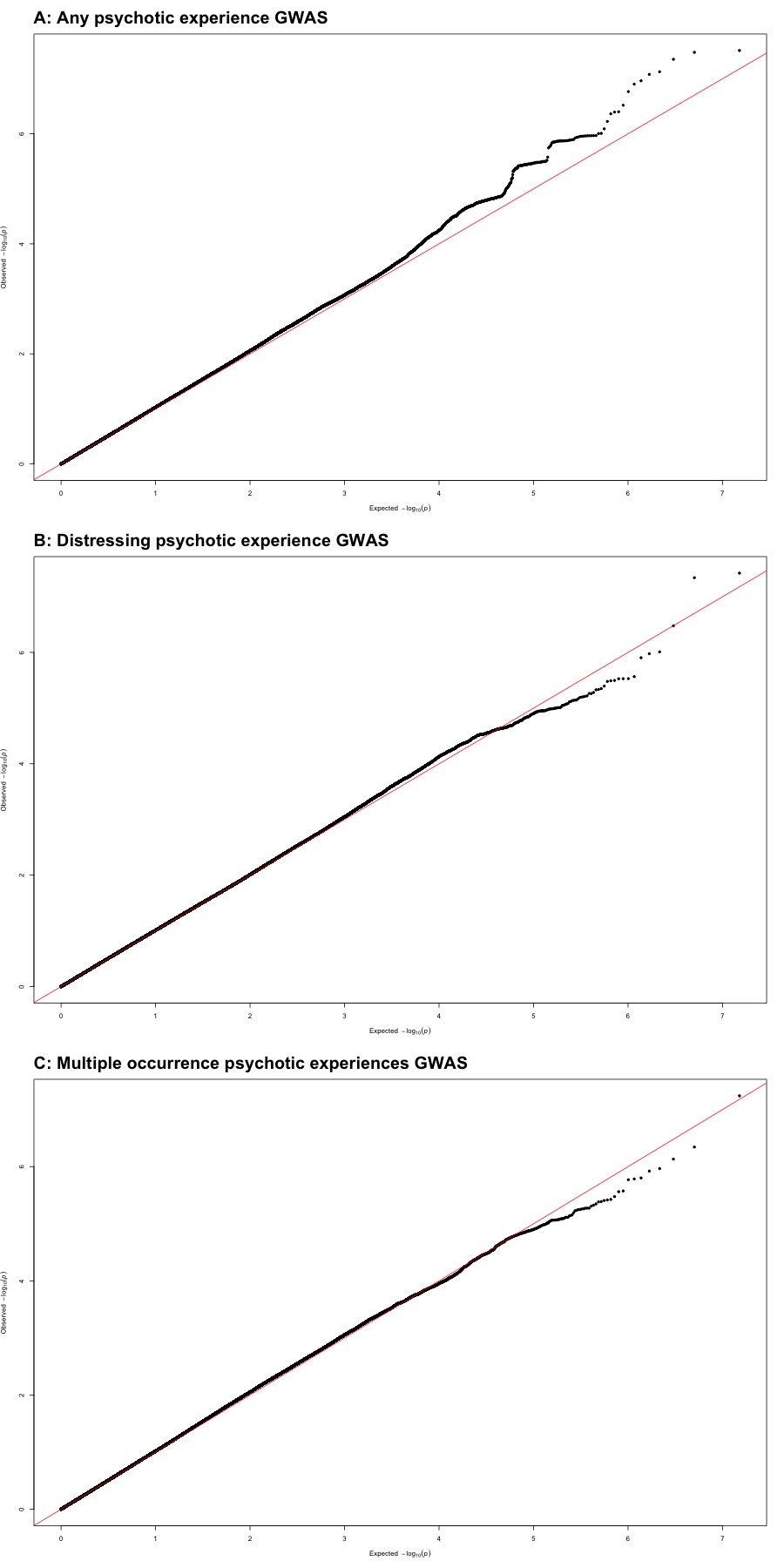

Quantile-quantile (QQ) plots of GWAS analyses. Plot A: any psychotic experience, plot B: distressing psychotic experiences, plot C: multiple occurrence psychotic experiences.

### Supplementary Figure 6: PRS analysis of PE symptoms in ALSPAC

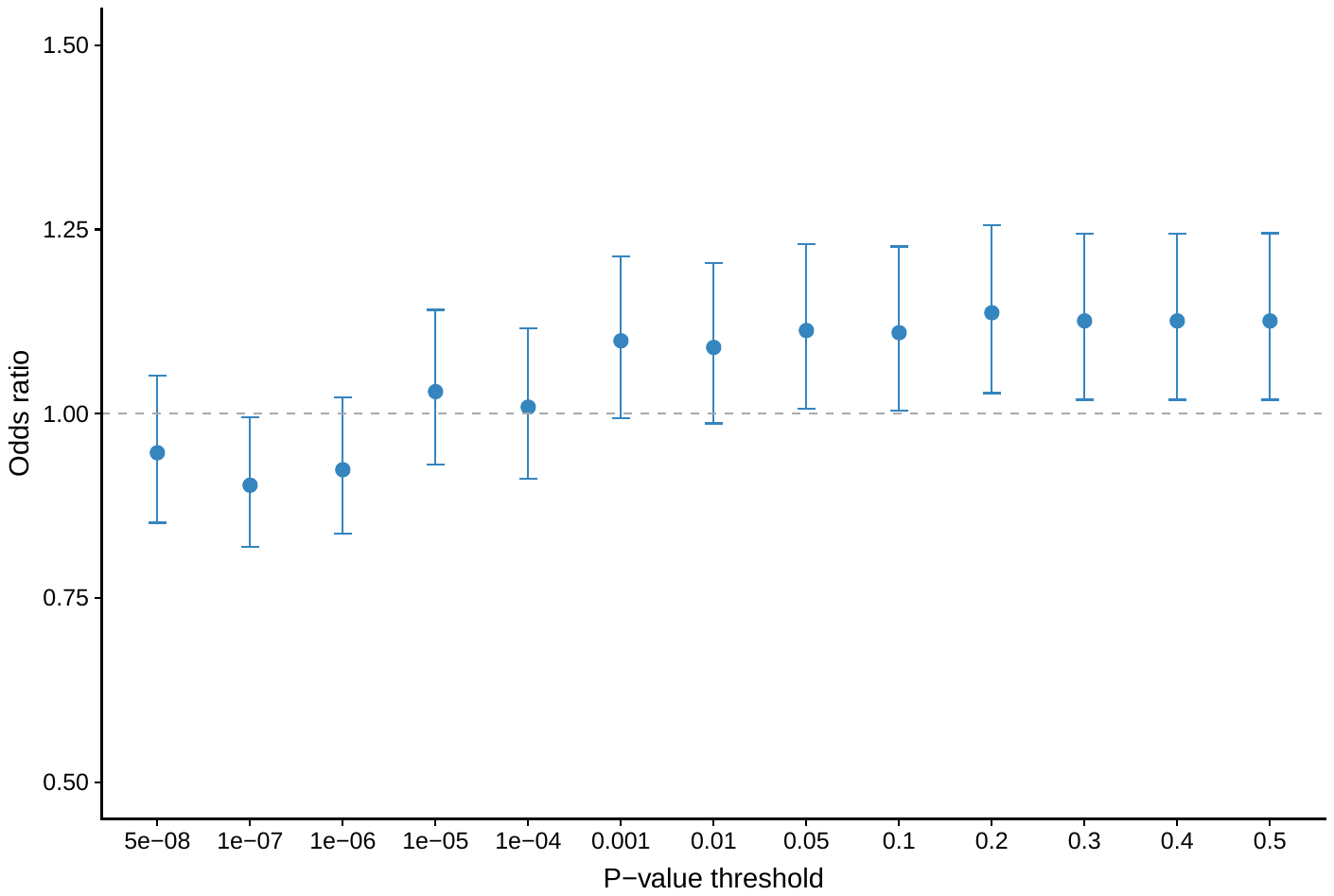

Polygenic risk score (PRS) analysis in ALSPAC cohort. PRS was created using summary statistics from the GWAS of any psychotic experience GWAS in UK Biobank and targeting psychotic experiences in ALSPAC (PLIKSi at 12 or 18, n = 5310, 7.76% with psychotic experiences).

### Supplementary Figure 7: PRS analysis of PE phenotypes

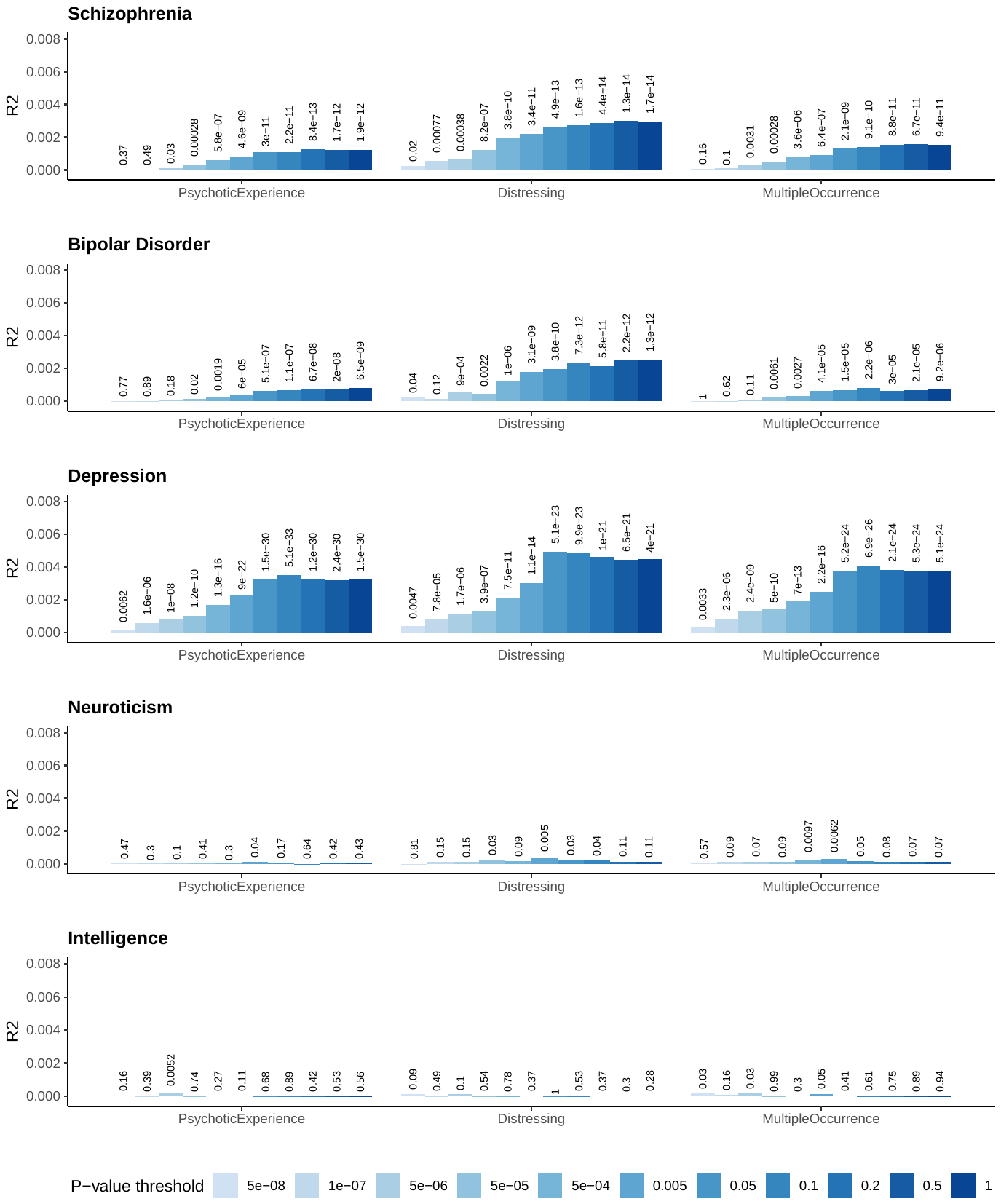

Polygenic risk score analysis. Each plot shows results for each PRS (schizophrenia, bipolar disorder, major depressive disorder, neuroticism and intelligence) and x axis shows psychotic experience phenotype. Bars represent variance explained (R2) and association p-values are listed above.

### Supplementary Figure 8: Average polygenic risk by PE phenotype

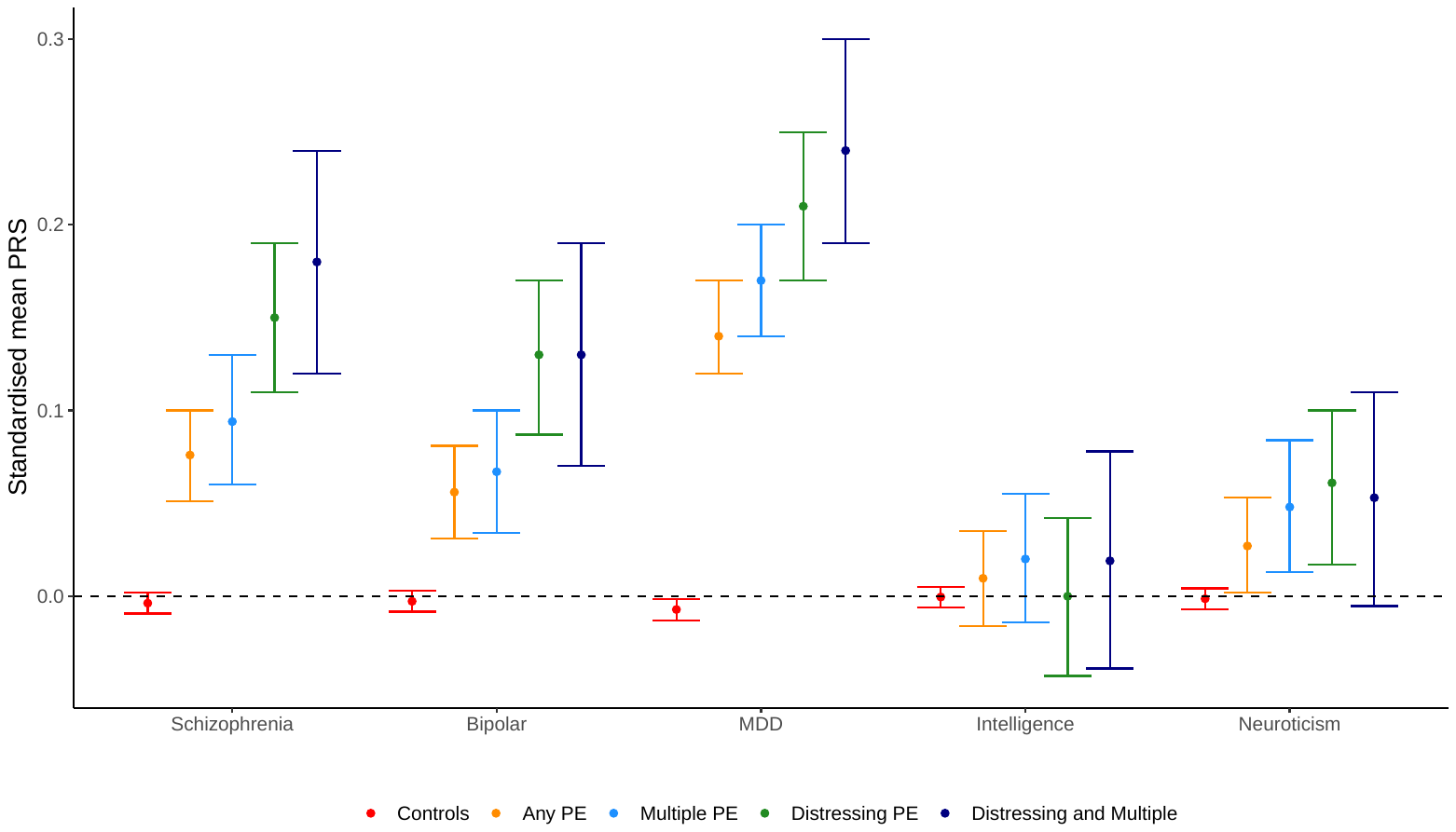

Mean polygenic risk score (PRS) for each psychotic experience phenotype.

### Supplementary Figure 9: PRS analysis of PE symptoms

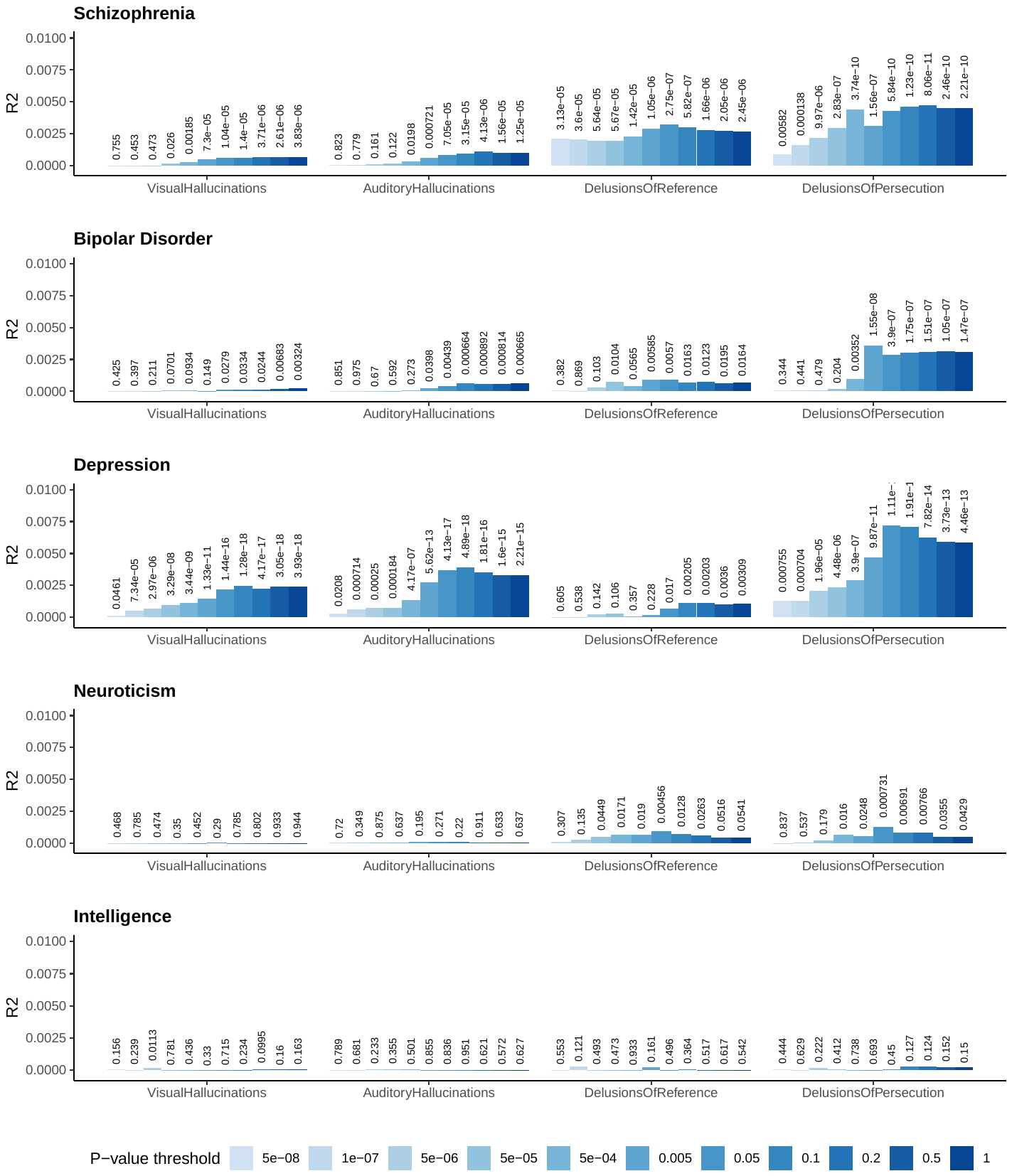

Polygenic risk score analysis. Each plot shows results for each PRS (schizophrenia, bipolar disorder, major depressive disorder, neuroticism and intelligence) and x axis shows psychotic experience phenotype. Bars represent variance explained (R2) and association p-values are listed above.

### Supplementary Figure 10: PRS analysis of PE symptoms

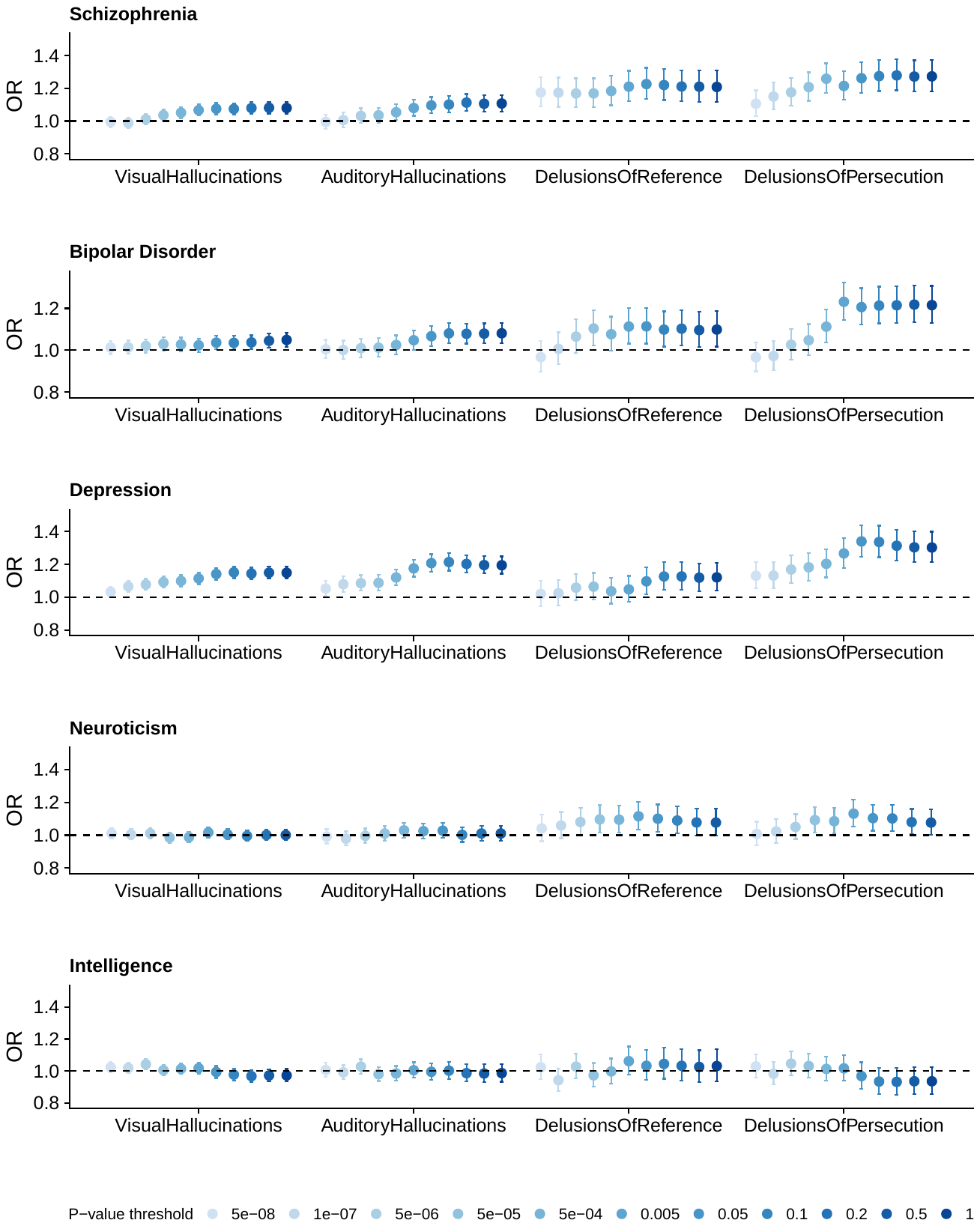

Polygenic risk score analysis. Each plot shows results for each PRS (schizophrenia, bipolar disorder, major depressive disorder, neuroticism and intelligence) and x axis shows psychotic experience phenotype. Bars represent odds ratio (OR) and 95% confidence intervals.
